# Bridging Auditory Perception and Natural Language Processing with Semantically informed Deep Neural Networks

**DOI:** 10.1101/2024.04.29.591634

**Authors:** Michele Esposito, Giancarlo Valente, Yenisel Plasencia-Calaña, Michel Dumontier, Bruno L. Giordano, Elia Formisano

## Abstract

Sound recognition is effortless for humans but poses a significant challenge for artificial hearing systems. Deep neural networks (DNNs), especially convolutional neural networks (CNNs), have recently surpassed traditional machine learning in sound classification. However, current DNNs map sounds to labels using binary categorical variables, neglecting the semantic relations between labels. Cognitive neuroscience research suggests that human listeners exploit such semantic information besides acoustic cues. Hence, our hypothesis is that incorporating semantic information improves DNN’s sound recognition performance, emulating human behavior. In our approach, sound recognition is framed as a regression problem, with CNNs trained to map spectrograms to continuous semantic representations from NLP models (Word2Vec, BERT, and CLAP text encoder). Two DNN types were trained: semDNN with continuous embeddings and catDNN with categorical labels, both with a dataset extracted from a collection of 388,211 sounds enriched with semantic descriptions. Evaluations across four external datasets, confirmed the superiority of semantic labeling from semDNN compared to catDNN, preserving higher-level relations. Importantly, an analysis of human similarity ratings for natural sounds, showed that semDNN approximated human listener behavior better than catDNN, other DNNs, and NLP models. Our work contributes to understanding the role of semantics in sound recognition, bridging the gap between artificial systems and human auditory perception.

## 1 Introduction

Recognizing sounds involves the conversion of acoustic waveforms into meaningful descriptions of the sound-producing sources and events. Automatic and effortless in humans, sound recognition poses a considerable challenge for artificial hearing. Various machine learning (ML) algorithms have been proposed that treat sound recognition as a classification problem. Typically, these algorithms entail the initial extraction of diverse features from the acoustic waveform, which are further analyzed and assigned to predefined classes. [1]. In recent developments, deep neural networks (DNNs) have emerged as superior to traditional ML algorithms in sound recognition tasks. Following parallel advancements observed in visual object recognition research, [2], convolutional neural networks (CNNs) have been used for sound classification tasks [3–5] (here, referred to as sound-to-event CNNs). Trained on a large-scale dataset of human-labeled sounds (Audioset, [6]), Google’s VGGish and Yamnet yield remarkable performance. These networks receive spectrogram representations as input and can classify sounds into a large number of classes (527 and 521 classes, for VGGish and Yamnet, respectively). Since their publication, VGGish and Yamnet (and related networks [5]) have been fine-tuned for applications in several specialized acoustic domains, from neonatal heart-beat and lung sound quality assessment [7] to aircraft detection system [8] and speech-emotion recognition [9].

In addition to the basic set of labels, Audioset [6] introduced a taxonomy specifying an additional set of super-ordinate labels and their (hierarchical) relations to the basic set. While DNN models frequently employ the Audioset basic labels (or subsets of them) for training purposes [10], the taxonomic information is generally not utilized (see [11] for an exception). This is because labels are commonly encoded as binary categorical variables using one-hot or multi-hot (in case of simultaneous multiple labels) encoding as depicted in Fig. 1. DNNs trained with this approach in fact map sounds to a set of orthogonal labels.

**Fig. 1.**
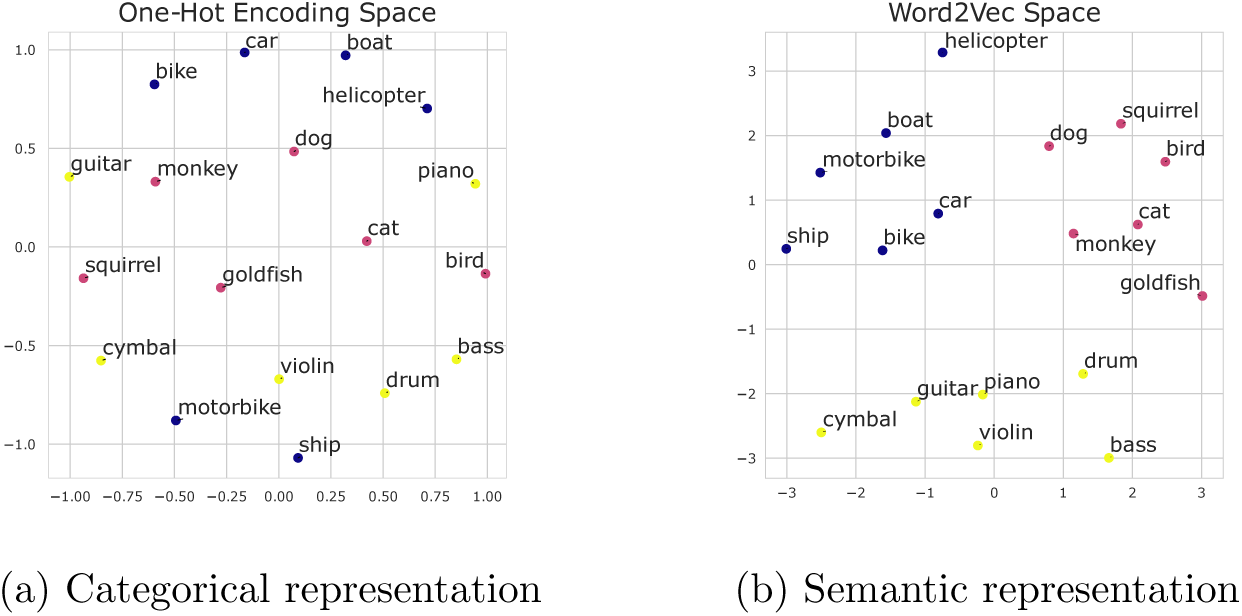
Categorical vs Semantic label encoding. Comparison of t-Stochastic Nearest Embedding (t-SNE)[21] visualizations between one-hot encoding (a) and Word2Vec (b) spaces: Embeddings were made by a one-hot encoding transformation of the words (a), or through the use of the GoogleNews-300D Word2Vec model[15] (b). In (a), words are equidistant from one another, and the proximity of words with semantic relationships follows the order in which the words are listed. However, in (b), words that are semantically related are closer to each other, demonstrating a more meaningful representation.

Research in cognitive psychology [12] and cognitive neuroscience [13] suggests that human listeners, when engaged in listening to and in comparing real-world sounds, exploit higher-level semantic information about sources in addition to acoustic cues. In a recent study by Giordano et al. [13], behavioral data involving perceived sound (dis)similarities, assessed through a hierarchical sorting task [14], were analyzed to investigate the explanatory power of sound-to-event DNNs, such as VGGish and Yamnet, and other models related to acoustic, auditory perception, and lexical-semantic (natural language processing, NLP). The results demonstrated that sound-to-event DNNs surpassed all other models in predicting human judgments of sound dissimilarity, indicating that sound-to-event DNNs provide, at present, the best approximation of human behavior for sound (dis)similarity judgments. In addition, the results highlighted the ability of NLP models, specifically Word2Vec [15] to capture variance in behavioral data that couldn’t be accounted for by sound-to-event DNNs trained with categorical labels.

Motivated by these findings, the present study sought to develop DNNs that - mimicking human behavior - incorporate lexical semantic information in the recognition of sounds. To this aim, we formulated sound recognition as a regression problem, training a convolutional DNN to learn the mapping of spectrograms to continuous and distributed semantic representations. In particular, we obtained these representations as the embeddings from NLP models. We considered word-level, pre-trained embeddings: *Word2Vec*[15], and context-dependent embeddings: Bidirectional Encoder Representations from Transformers (*BERT*) [16]. Additionally, we considered the semantic embeddings obtained from the Contrastive Language-Audio Pretraining (*CLAP*) text encoder, a contrastive-learning model that brings audio and NLP BERT embeddings into a joint multimodal space [17].

To evaluate the impact of semantics on sound recognition, we trained two types of DNNs: semDNN, utilizing one of the described continuous semantic embeddings, and catDNN, employing categorical, one-hot encoded labels. To ensure a fair comparison, we trained the DNNs from scratch using a curated dataset of 388,211 sounds from the Super Hard Drive Combo [18]. In this dataset, a rich semantic description of each sound can be derived from the associated metadata. We expected that, compared with a homologous network trained with categorical labels, semDNN would produce semantically more accurate labeling in sound recognition tasks and that semDNN embeddings would preserve higher-level lexical semantic relations between sound sources. Furthermore, we expected that semDNNs would better approximate human behavior in auditory cognitive tasks compared to catDNNs due to the preservation of semantic relations in NLP embeddings. Our approach differs from previous studies that combined sound-to-event DNNs with language embeddings [17, 19], as we specifically focus on evaluating the effects of semantic representation types and predicting human perceptions. In summary, our work aims to bridge the gap between artificial sound recognition systems and human auditory perception by incorporating semantic information into DNNs [20].

## 2 Methods

In this section, we outline the methods used in our study (framework depicted in Fig 2). We describe our strategy for label encoding using semantic models and how it differs from one-hot encoding. We address the limitations of context-dependent models for single-word embeddings, and the motivation for dimensionality reduction for comparing different models. We then detail the network architecture, the training dataset, and the strategy adopted to limit the influence of class unbalancing on model training. We subsequently evaluate our models using four different external datasets in addition to the internal one, describe the word retrieval strategy to get labels from the embeddings, and discuss the evaluation metrics adopted to assess the models’ performances. Finally, we provide a summary of a cross-validated Representation Similarity Analysis (RSA) carried out to predict human judgments of sound dissimilarity and provide a description of the compared models.

**Fig. 2.**
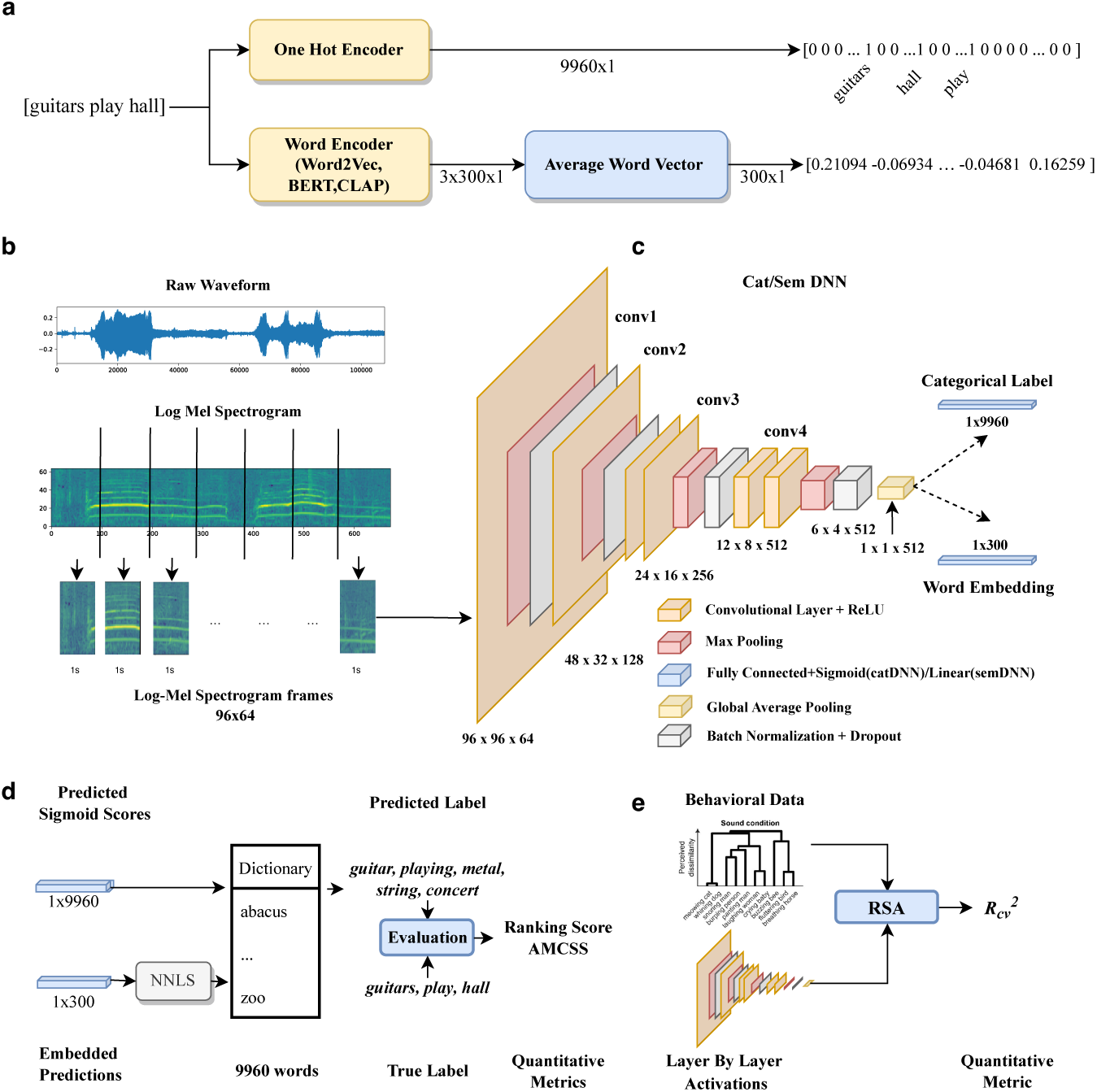
Proposed framework: from preprocessing to evaluation. **a)** Label encoding strategy: transition from lexical units to either orthogonal (one-hot encoder) representations or continuous representations (word encoder). **b)** Audio Preprocessing: conversion of waveforms to 1 s patched spectrograms. **c)** Model Architecture: Note the variation in the final dense layer. **d)** Evaluation Phase: transition from embeddings to word-level predictions, with the computation of quantitative metrics to gauge model performance. **e)** Model Comparison: Representational Similarity Analysis (RSA) is adopted to compare the ability of models to predict human behavioral data.

### 2.1 Semantic Models

We employed three language models for label transformation: GoogleNews Word2Vec-300D [15], BERT-768D [16], and CLAP-1024D text encoder [17].

Word2Vec is a word-based encoder trained on large corpora to learn distributed representations that capture semantic similarities and relationships between words.

BERT, Bidirectional Encoder Representations from Transformers, is a pre-trained language model that learns to capture deep relationships and context between words in sentences.

Contrastive Language-Audio Pretraining (CLAP) is a transformer-based architecture that is fine-tuned for the audio-to-text task using a large dataset of paired audio and text descriptions encoded with BERT. The text encoder is trained jointly with the audio encoder using a contrastive loss function, which encourages the audio and text representations to be similar in the joint multimodal space. Specifically, the contrastive loss function aims to maximize the similarity between the representations of a given audio-text pair while minimizing the similarity between the representations of different pairs.

#### 2.1.1 Label Encoding Strategies

To extract semantic embeddings from the sound descriptions we performed the label transformation depicted in Figure 2, panel a). For CatDNN, we used one-hot encoding. Each label was represented as a binary vector of 9960 dimensions, as the number of entities contained in the dictionary (see 2.3), with a value of 1 indicating the presence of the label in the description.

To obtain a single embedding describing the sound semantics in SemDNN, we directly used the single-word labels as input to word-based encoders. Specifically, we computed the Word2Vec, BERT or CLAP embeddings for each word present in the label and then averaged these word embeddings. This resulted in a single embedding that captured the overall semantic information of the sound (sound-level embedding). This process was straightforward for Word2Vec, as it produces a single, context-independent embedding for each word. However, for BERT and CLAP, we needed to make some preliminary adjustments before applying the same method.

##### BERT and CLAP embeddings

BERT is a context-dependent language model, which means that the embedding of a single word changes depending on its position in the sentence, the surrounding words, and the sentence length. To obtain word-level BERT embeddings, we first considered all the sentences contained in the SoundIdeas dataset (see section 2.3). From these sentences, we generated an initial dictionary consisting of the words present in the sentences. This dictionary was specifically designed for BERT representation and associated each word with two elements: its single-word embedding within a particular sentence and the corresponding sentence itself. This approach was taken in order to capture the variations in single-word BERT embeddings across different sentences where the word appears. The computation of these sentence-dependent word-level BERT embeddings required careful handling of tokenization. For this, we utilized the *bert-base-uncased* model and its built-in tokenizer. It is worth noting that, when calculating the BERT embeddings for single words, we focused on the word-specific token representation. This strategy differs from using the [CLS] token, which represents the entire sentence’s embedding. The reason behind this decision was to ensure a more granular representation of individual words. In contrast, the [CLS] token, although it represents the overall semantic content of the sentence [16], does not provide a focused representation of each unique word within the sentence. We then averaged the embeddings associated with each word across different sentences to obtain a final word-level BERT embedding. This averaging was motivated from the fact that sentence-dependent word-level BERT embeddings are more similar among them compared to embeddings of different words. This is illustrated in Figure 3a for 10 sampled words from the dictionary. For these words, we calculated the cosine similarity between pairs of vectors reflecting context-sensitive word embeddings. This resulted in a similarity matrix, which is visualized as a heatmap, where brighter squares indicate higher similarity and darker squares indicate lower similarity. It can be observed that sentence-dependent BERT embeddings exhibit contextual variations, but are still more similar among them than to the other words. Thus, averaging across sentences allowed us to obtain contextually robust word-level embedding for each word and reduce the BERT dictionary to the same dictionary we used for Word2Vec.

**Fig. 3.**
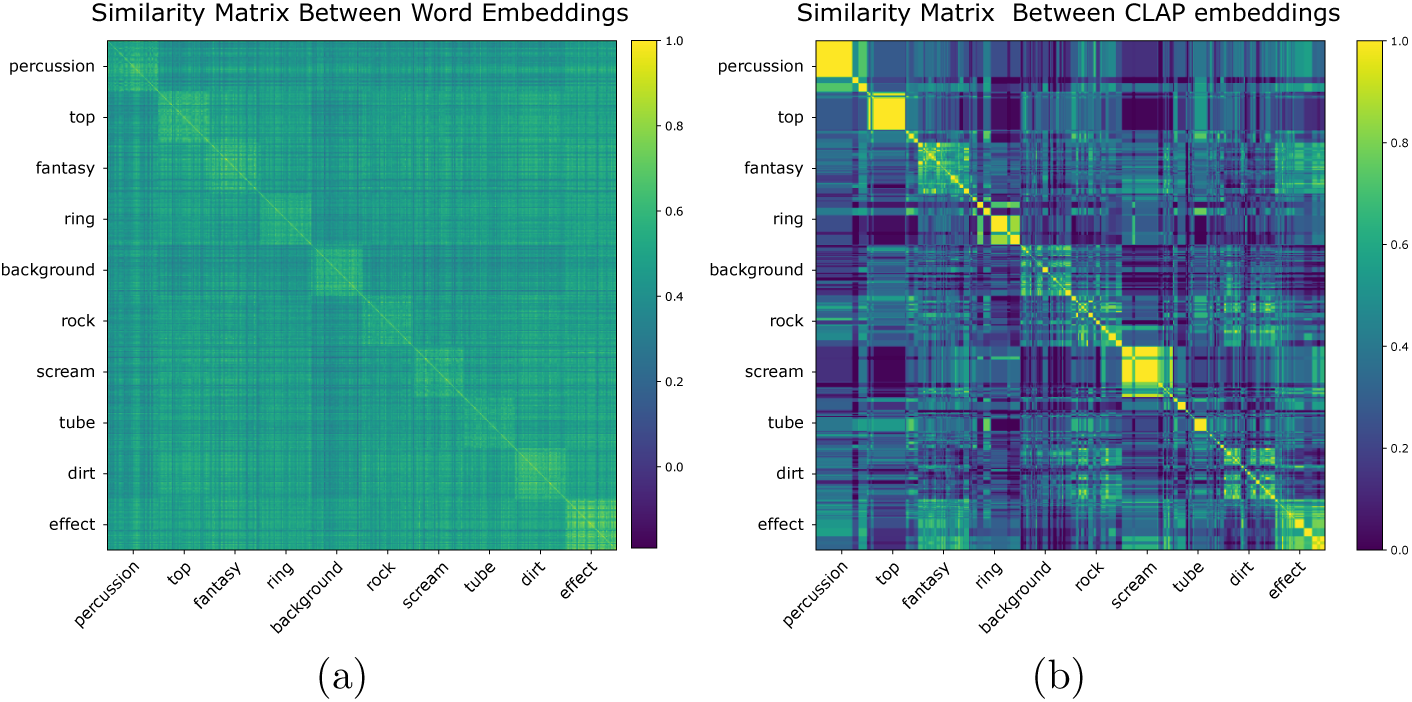
BERT and CLAP normalized Similarity matrices. Comparing BERT and CLAP embeddings across 100 different sentences reveals interesting patterns. While BERT embeddings exhibit noticeable variation for each word, implying some degree of divergence, CLAP embeddings display remarkable consistency and reduced variation.

Unlike BERT, CLAP is fine-tuned to reduce dissimilarity between audio and text pairs in a multimodal setting. As part of this process, CLAP aligns audio and text representations to occupy a joint multimodal space [17]. This alignment ensures that similar audio and text pairs are closer together, while dissimilar pairs are farther apart. During the fine-tuning process, the model weights, including those used to generate embeddings, are updated to minimize the loss on the specific task, which is a symmetric cross-entropy loss function. Furthermore, CLAP generates a single embedding per sentence or word because it is a fine-tuned version of BERT, with an additional Dense layer at the end of its architecture. As a result, CLAP embeddings exhibit less variation compared to BERT embeddings. This can be observed in Figure 3b. Given these reasons, there is no need to compute multiple-word embeddings per word across different sentences. Therefore, we constructed a dictionary where each word contained in the Super Hard Drive Combo’s labels is associated with its word embedding generated by the CLAP-text encoder.

BERT and CLAP embeddings have a dimensionality of 768 and 1024, respectively, whereas Word2vec embeddings have 300 dimensions. Thus, as a final step for calculating, we reduced the dimensionality of BERT and CLAP embeddings using an autoencoder (see supplementary material, **??**). This reduction brought the embeddings to the same dimensionality as the Word2Vec model (300), while resulting in a negligible infromation loss (autoencoder reconstruction loss was 0.89% for BERT and 0.005% for CLAP,see suppl. material). We then substitute the original length embeddings of the preliminary BERT and CLAP dictionaries with theses reduced ones.

Thus, by calculating word-level embeddings and aligning the dimensions of all the models, we ensured a fair and meaningful comparison across the different semantic models. All the dictionaries have the same 9960 entities, extracted from the SoundIdeas sound labels, and an associated word representative 300-dimensional embedding. Sound-level embeddings are then obtained as the average of all word-level embeddings in the sound description.

### 2.2 Network Architecture

We developed two different neural network configurations for sound recognition task: semDNN and catDNN (Fig. 2c). Both networks resemble the VGGish [3] architecture and share similar components, such as four main convolutional blocks with 64, 128, 256, and 512 filters. Compared to VGGish [3], we added a dropout layer (rate = 0.2; [22]) and a batch normalization layer [23] after each down-sampling operation, and after the fully connected layers to improve the model’s generalization ability, prevent overfitting, and facilitate more stable and efficient training in comparison to VGGish. We also applied global average pooling after the last convolutional block to summarize the feature maps into a fixed-length vector. However, they differ in the output layer. Whereas VGGish has a 128-unit dense layer, SemDNN has a 300-unit layer with linear activation, and catDNN has a 9,960-unit dense layer with a sigmoid activation function. We used a different loss function for each of these architectures. For semDNN, we used an angular distance loss function, due to the nature of the regression task that aims to minimize the angle between the true word embedding and the word embedding predicted during training. This loss function is suitable for semantic embeddings, as it encourages the network to learn the continuous representation of words within the fitting domain.[15, 24–26]. On the other hand, catDNN uses a binary cross-entropy loss function, which is suitable for the multi-label classification task [27]. This loss function measures the difference between the predicted probabilities and the true labels and encourages the network to learn a discrete representation of words that can be used for classification.

#### SemDNN and its variant

We employed different strategies to train SemDNN. Specifically, we trained SemDNN using the Word2Vec, BERT, and CLAP representations as labels. Furthermore, as a purely acoustic approach, we trained a Convolutional Auto Encoder (CAE) with the architecture depicted in Figure 2c for the encoder, and a reversed architecture for the decoder. The CAE was trained using only acoustic inputs, without involving a categorical/semantic label. The Mean Square Error was employed as the loss function for the CAE. To provide an additional control network, we also considered SemDNN with random Normal-HE initialization [28], without training it. A summary of the variants is depicted in Table 1. Additionally, to assess the efficacy of semantically balanced training, we trained SemDNN using a randomly chosen dataset of the same length as the training set that was generated from hierarchical clustering (see section 2.3).

**Table 1.** Summary of SemDNN Variant Networks.

| Network | Key Aspects |
| --- | --- |
| **SemDNN** | Utilized Word2Vec [15] representation as labels |
| **SemDNN<sub>BERT</sub>** | Utilized BERT [16] representation as labels |
| **SemDNN<sub>CLAP</sub>** | Utilized CLAP [17] representation as labels |
| **SemDNN<sub>unbal</sub>** | Trained with randomly chosen training dataset |
| **SemDNN<sub>noTrain</sub>** | No training, random Normal-HE[28] initialization |
| **CAE** | Mirrored encoder architecture of SemDNN, see Fig 2 (c) |

#### Preprocessing and Input Features

The input audio clips were preprocessed as follows: first, we resampled signals to a standard 16 kHz sampling rate format and converted them to mono. Then, we split the clips into non-overlapping segments of 960 ms. For each segment, we computed a short-time Fourier transform on 25 ms Hanning-windowed frames with a step size of 10 milliseconds. This allowed us to break down the signal into its constituent frequencies at each moment in time and perform a detailed analysis of the audio data. Next, we aggregated the resulting power spectrogram into 64 mel bands covering the range of 125-7500 Hz.

Finally, we generated a stabilized spectrogram consisting of 96-time windows per 64 log mel bins. To obtain the log mel spectrogram, we took the logarithm of the mel spectrogram values. Additionally, we applied a stabilization technique to prevent numerical instability during this step. The stabilization involved adding a small offset of 0.01 to the mel-spectrum before taking the logarithm. This offset ensures that the logarithm operation does not encounter zero values, which could lead to undefined or erroneous results. The resulting stabilized spectrogram was then utilized as the input for training and evaluation of the deep neural networks (DNNs) and it is the same procedure applied in [3]. Each 1 s sound frame inherited the same label.

### 2.3 Training Dataset

The networks have been trained using sounds and labels from SuperHard Drive Combo (SHDC) by Sound Ideas [18], a collection of 388,199 variable-length sounds (2,584 hrs) covering a wide range of sound sources and events. SHDC contains 7 different natural sound databases that can be considered as independent datasets: DigiEffects [29], General Hard Drive Combo [30], Hollywood Edge [31], Mike McDonough Speciality [32], Serafine[33], SoundStorm [34], and Ultimate [35].

We employed a natural language processing (NLP) pipeline to extract a dictionary of sound-descriptive words from the SHDC metadata. The initial step involved eliminating all non-informative tokens from the filename metadata. This included numbers, serial IDs, stop-words, and all non-English words that were not included in the GoogleNews300D-Word2Vec[36] model dictionary. In the majority of cases, these filenames contained information about the sound sources and events occurring in the sounds. For instance, “*ManSneezesWhugeLoCRT* 026004*.wav*” was reduced to *“man sneezes”*, and “*Rhythmic − Percussion − V ariation − Short − V ersion −* 21*PET* 10 *−* 088*.wav*” was transformed into *“rhytmic percussion variation”*. Our next step was to replace nouns that were either too specific (subordinate categories) or too general (super-ordinate categories) with basic-level descriptors. For example, in first case, specific car models like *“Subaru Impreza”* or *“Audi TT”* were replaced with the more general term *“car”*, and specific dog breeds like *“labrador”* or *“pincher”* were replaced with *“dog”*. In the rare case of super-ordinate categories, expressions that were excessively vague such as “animals” were replaced with basic-level descriptors that provided more specific information. For instance, for the file that was called *“AnimalV arious DIGIMEGADISC −* 60*.wav”*, we replaced *“ani-mal various”*, after listening to the sound with *“lion growls bats swarm”*, thus preserving the semantic integrity of the sound while avoiding excessive generality. This process was initially automated using the NLP pipeline. To ensure accuracy, the results were manually reviewed and corrected as necessary, as demonstrated in the example mentioned above. The decision to standardize descriptors to a basic level was driven by the need to balance specificity and generality, maintaining meaningful semantic information without overloading the model with excessive detail. This approach allows for a more manageable and semantically consistent representation of heterogeneous natural sounds, enhancing the model’s ability to learn and generalize from the sound-descriptive words in the SHDC metadata [37]. The resulting output of the aforementioned NLP pipeline was sound labels extracted from the filenames of the sounds and an entities-dictionary of 9960 units. The first two columns of table 2 show some examples, note in the last row how the transformation in the base-level category occurs, from *Harrier* to *jet*.

**Table 2.**
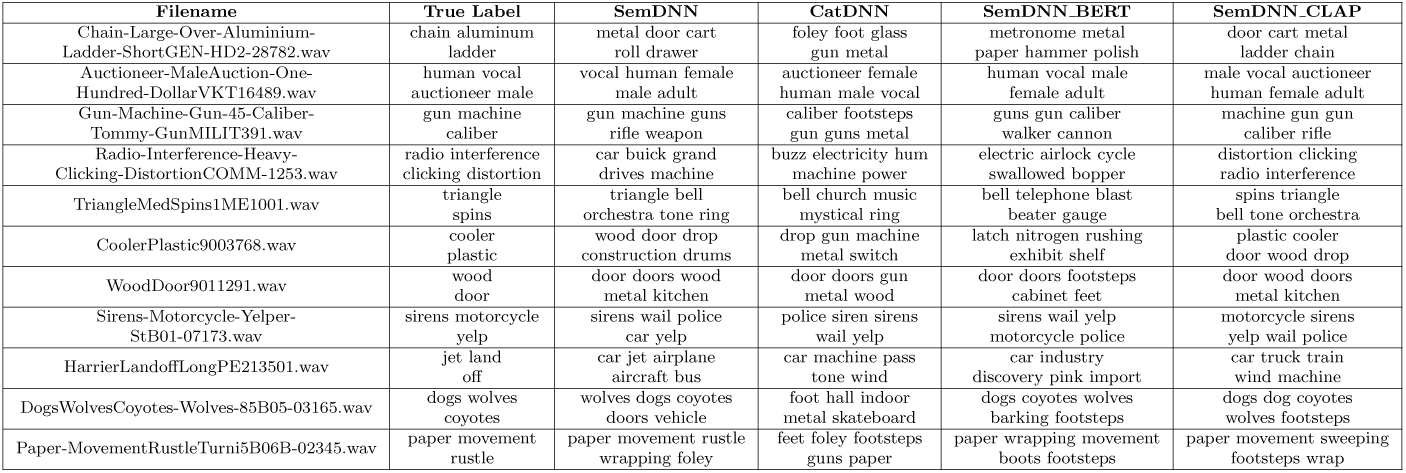
Top 5 Predictions. Top-5 word predicted from NNLS (for SemDNN and its variants) and sigmoid activations (for CatDNN). Note also the results of the NLP pipeline to retrieve labels from the sounds’ filename.

An initial analysis of the word-frequency distribution in the database (Fig. 4) revealed that sound labels were highly skewed towards words such as “car”, “door”, “metal”, and “engine” from a prominent portion of the database dedicated to vehicle sounds. To rectify this imbalance, we implemented a semantics balancing procedure relying on a hierarchical clustering analysis of the Word2Vec embeddings of the sound-descriptors dictionary. We initially computed the Word2Vec embedding of each word and generated a normalized pairwise cosine similarity matrix. This matrix was subsequently input to a hierarchical clustering algorithm (ward-linkage [38]). Different cluster counts (100, 200, 300, 400, 500) were tested to assess the impact of clustering granularity on the performance, which was measured using the evaluation procedure described in the section 2.5. Our results indicated that the optimal performance was obtained with 300 clusters (see Figure **??** in the Supplementary Material). Finally, we randomly selected up to 20 words from each cluster, matching the average number of words per cluster. We also chose 300 sounds for each of the selected words, leading to a more balanced dataset. The resulting balanced dataset included 273,940 sounds (training set = 90% = 246,546 sounds; 1,366,848 frames; validation set 5%; internal evaluation set = 5%).

**Fig. 4.**
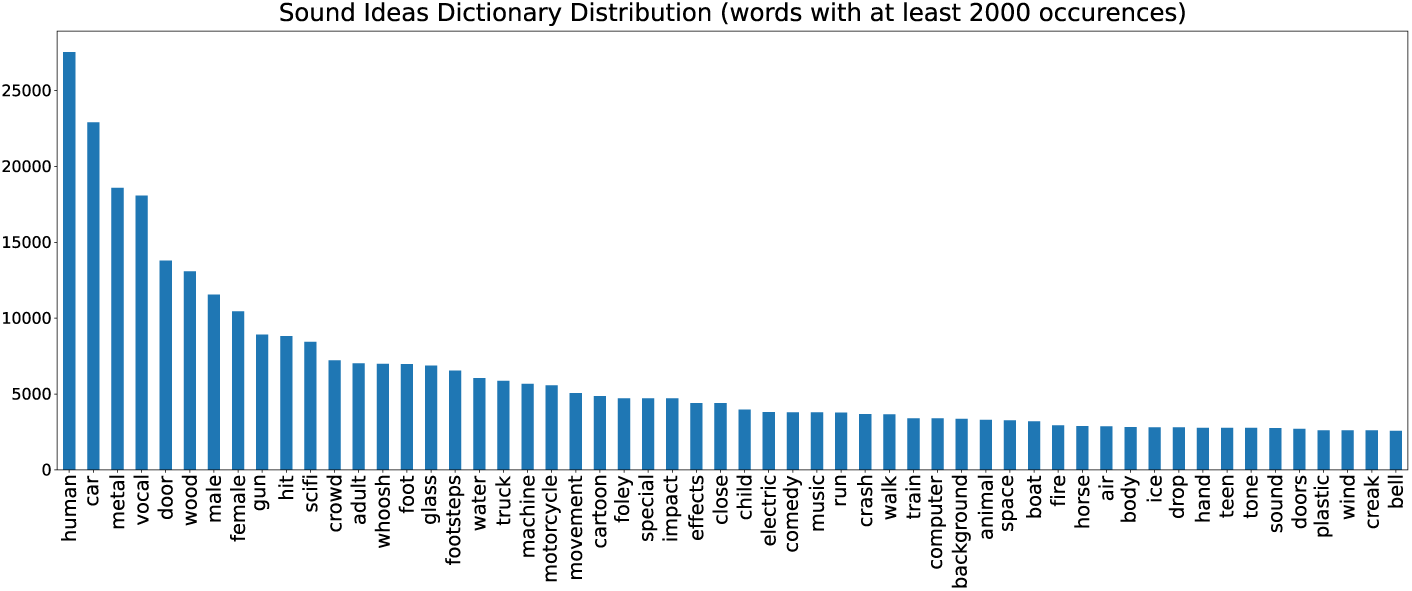
Labels distributions, a: The most frequent word in the dataset is “car”, followed by “door” and “metal”. To avoid over-representation of the most frequent classes, we developed and applied a method for creating a semantically-balanced dataset.

In the next phase, we conducted a quality check to evaluate the spatial arrangement of the words within clusters in the embedding space. Specifically, we ranked the clusters based on their inter-cluster cosine similarity, from highest to lowest. Utilizing t-Stochastic Nearest Embedding (tSNE) [21], we visualized the top 25 clusters, ensuring the words within each cluster were semantically related (see Fig. 5). The figure results in a visual representation of the semantic space. Each point in the plot corresponds to a semantically related word contained in a specific color-coded cluster. The top 25 clusters are highlighted, showcasing the arrangement of words in the reduced-dimensional space.

**Fig. 5.**
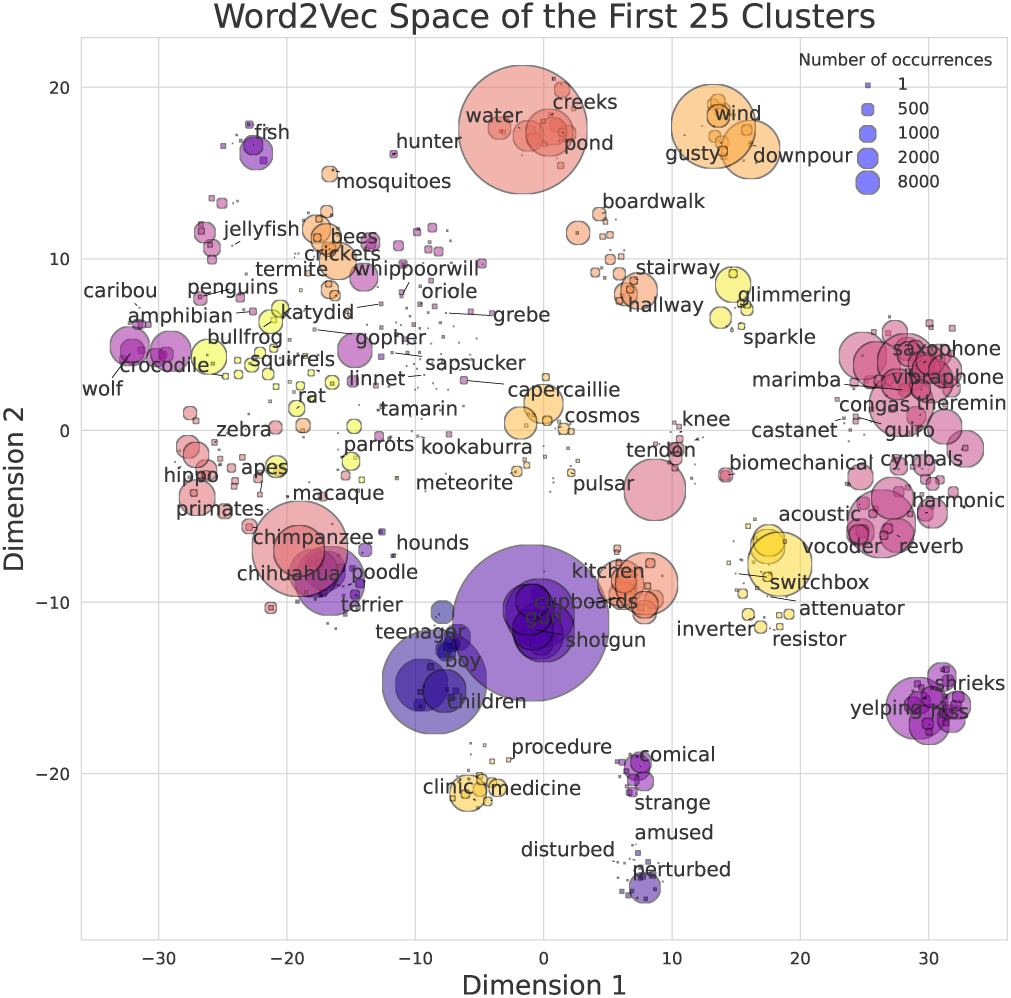
t-SNE visualization of the top 25 semantically related word clusters. Spatial arrangement of words in the embedding space, where each point represents a semantically related word. Color-coded clusters highlight the organization of words in the reduced-dimensional space, providing insight into the ideal relationships between spectrograms and their associated semantic representations.

### 2.4 Evaluation Datasets

We evaluated the performance of our proposed approach with four publicly-available natural sound datasets: FSD50k [10], consisting of 10,231 44.1 kHz mono audio files and 200 labels; Environmental Sound Classification-50 (ESC-50) [39], made up of 2,000 5-second, 44.1 kHz mono, sounds and 50 label-classes; Urban Sound 8K [40], comprising 8,732 sounds with lengths of up to 4 seconds, 44.1 kHz mono, and 10 class labels; and Making Sense Of Sounds [41], which includes 500 5-second, 44.1 kHz mono, sounds divided into two level categories, 5 macro-classes, and 91 subclasses. In addition, we used a 5% subset of the SoundIdeas dataset consisting of 13,697 sounds (not used for training our models) to evaluate the performance of our models (internal evaluation).

### 2.5 Semantic-learning accuracy

We compared semDNN and catDNN using two prediction-accuracy metrics: Ranking score and Average Max Cosine Similarity (AMCSS). For the different variants of semDNN, which produce semantic embeddings as predictions, we employed non-negative least squares (NNLS) regression [42] to convert the embeddings back into word predictions. The models’ training involved generating embeddings using Word2Vec, BERT, and CLAP for individual labels in the sound descriptions. However, in the evaluation phase (see 2.5) these embeddings were averaged to create one single representative semantic embedding for each sound. To retrieve the constituent single-word embeddings from the predicted mixture, we used a Non-Negative Least Squares (NNLS) [42] approach. The NNLS regression projected the predicted semantic embeddings onto the single-word embedding space, considering the entire dictionary as the design matrix of dimension 9960×300. By applying a non-negativity constraint in the NNLS, the coefficients of the linear combination remained non-negative, preserving the original averaging process.

#### Ranking Score

To evaluate the prediction accuracy of the NNLS regression coefficients, known as *β*-values (for semDNN, in all its variants) and the sigmoid output probabilities (for catDNN), we employed a ranking-based metric called the “ranking score.” This metric allows us to compare the models’ predictive abilities while considering the relative positions of the true labels within the sorted predictions. First, we obtained the NNLS *β*-values, which represent the coefficients assigned to the different words of the dictionary (9960 *β*-values). The obtained *β*-values were sorted in descending order based on their magnitudes.

Similarly, we obtained sigmoid output probabilities from CatDNN and we sorted with the same criteria. These probabilities represent the model’s confidence scores for each possible class or label. To calculate the ranking score, we utilized the sorted predictions. The ranking score is defined as follows:

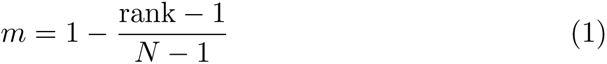

Here, *m* represents the ranking score, *N* represents the length of the dictionary, and rank is the position in the dictionary of the predicted label corresponding to the true label. We computed the ranking score individually for each word in multi-word labels and then averaged the scores.

The ranking score penalizes predictions that deviate significantly from the true labels, resulting in a lower score for predictions ranked further away from the true label. Conversely, a higher score indicates a closer alignment between the predicted label and the ground truth. The ranking score is threshold-independent, allowing a comprehensive comparison of all words in the dictionary (9960 words) with their respective true labels.

#### Average Maximum Cosine Similarity Score (AMCSS)

To compare the performance of the different networks we used a novel metric, the Average Maximum Cosine Similarity Score (AMCSS). The AMCSS (Average Maximum Cosine Similarity Score) is computed by considering the predicted labels and true labels. The true labels are extracted from a fixed dictionary, which is described in Section 2.1.1. The AMCSS is defined as:

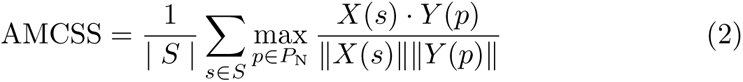

where *| S |* represents the number of true sound labels, *S* represents the set of true sound labels, *P* represents the set of predicted labels, *P*_N_ represents the top *N* = 10 words obtained from the predicted labels, *X*(*s*) represents the word embeddings of the true sound label *s*, *Y* (*p*) represents the word embeddings of the predicted label *p* and the operation in fraction calculates the cosine similarity between the word embeddings *X*(*s*) and *Y* (*p*).

To calculate the AMCSS, we compare the word embeddings of the true labels and the top 10 words obtained from the NNLS (for Word2Vec, BERT, and CLAP) or sigmoid output (for catDNN) generated from the model predictions. We calculate the cosine similarity between each word embedding in the true labels and the top 10 words. The maximum cosine similarity value among all the comparisons is taken as the AMCSS. The AMCSS is computed as the average of the maximum cosine similarity scores for each word, ensuring more robustness as compared to using only the top word from NNLS (or sigmoid). The AMCSS reflects the network’s ability to identify relevant words and concepts associated with the true label, even if the exact label is not among the top predictions.

However, such a metric is influenced by the geometry of the manifold where the embeddings lie, and it is therefore misleading to directly compare the AMCSS obtained with Word2Vec with the one obtained, for instance, with BERT. We illustrate this problem more in detail in Fig. 6, where we considered 500 randomly chosen sounds from the internal test set and computed the cosine similarity matrix between the predicted and true embeddings of all the sounds. The upper row represents the values on the main diagonal of the similarity matrix, *i.e.* the cosine similarity between the sound embedding and its prediction, for each sound. The lower row displays instead the values outside the main diagonal, thus the cosine similarities between the prediction of a sound embedding, and the true embeddings of different sounds. A model that discriminates well a correct predictions would result in high values for the diagonal elements (top row), and lower values for the off-diagonal elements (lower row). Notably, BERT and CLAP exhibit high cosine similarity values both on the diagonal and off-diagonal, resulting in a right-skewed and lower variance distribution. In addition, the difference between the mean values of the diagonal and off-diagonal is considerably smaller for BERT and CLAP. On the other hand, Word2Vec, despite having lower overall similarity values, demonstrates higher selectivity, showing greater differences between the mean values of diagonal and off-diagonal elements. Based on these findings, we computed AMCSS using the same dictionary for all language models and decided to use Word2Vec as a reference, ensuring a more discriminative metric to compare models.

**Fig. 6.**
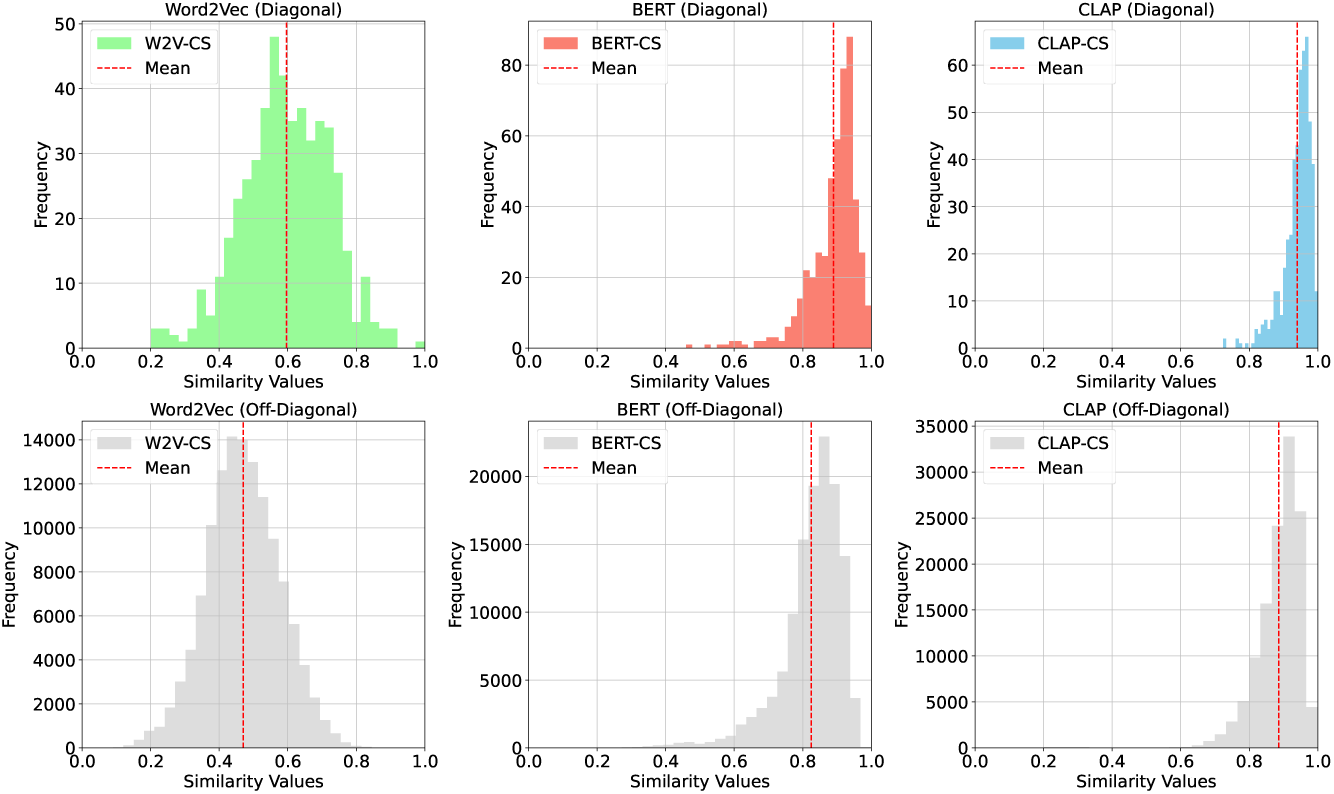
Distribution of Cosine Similarities between True and Predicted Embeddings for Different SemDNN Variants. Cosine Similarity (CS) distributions between true embeddings and predictions of 500 random sounds from the SoundIdeas [43] test set. The upper row represents the cosine similarity between the sound embedding and its prediction, measuring the network’s accuracy in predicting embeddings for the same sound (diagonal values of the similarity matrix). The lower row shows the cosine similarities between the prediction of a sound embedding, and the true embeddings of different sounds (off-diagonal values of the similarity matrix), reflecting the network’s performance in comparing a reference sound to different sounds.

### 2.6 Behavior prediction accuracy

We evaluated to what extent layer-by-layer embeddings of semDNN and its variants and catDNN, and of several control networks, including the CAE, predicted perceived dissimilarity judgments obtained with humans.

#### 2.6.1 Behavioral Data

In Giordano et al.’s study (Experiment 2, [14]), data were collected from two groups, each with 20 participants. Random assignment placed participants in either the sound dissimilarity or word dissimilarity condition. In the sound dissimilarity condition, participants estimated the dissimilarity between 80 natural sounds. In the word dissimilarity condition, participants assessed the dissimilarity of sentences describing the source of each sound (e.g., “meowing cat”). For the behavioral datasets, name plus verb sound descriptors were derived from the results of a preliminary verbal identification experiment ( [14], Experiment 1), during which 20 individuals, who did not take part in Experiment 2, were asked to identify the sound-generating events using one verb and one or two nouns. In particular, for each of the sound stimuli, the name plus verb sound descriptors considered for the analyses in this study, and evaluated by participants in the word condition, were the modal verbs and nouns (that is, the most frequent verbs and nouns) across the 20 participants in the verbal identification experiment. Each condition involved evaluating two sets of 40 stimuli categorized as living or non-living objects. The stimuli had a median duration of 5.1 seconds. Sessions were conducted separately for each stimulus set, with the presentation order balanced across participants. Participants performed a hierarchical sorting task. Initially, they grouped similar sounds or verbal descriptors into 15 groups using onscreen icons. Clicking on the icons activated the corresponding stimuli. Participants then iteratively merged the two most similar groups until all stimuli were consolidated into one group. The dissimilarity between stimuli was determined based on the merging step at which they were grouped, with dissimilar sounds or words being merged at a later stage of the procedure than similar sounds or words. The resulting output is a dissimilarity matrix.

#### 2.6.2 Cross-validated Representation Similarity Analysis

We employed a cross-validated computational modeling framework, similar to Giordano et al. [13], to predict behavioral dissimilarities using model distances derived from the network representations. To this purpose, we initially computed cosine the distance between stimuli within each layer of a specific network (encoder-only for CAE). For each network separately, we then used layer-specific distances to predict group-averaged behavioral dissimilarities within a cross-validated linear regression framework. More specifically, we adopted a repeated 10-fold cross-validation split-half approach to estimate the behavior variance 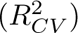 predicted by each network (100 random splits of participants into training and test groups, with independent standardization of group-averaged training and test dissimilarities). 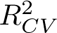 was estimated as 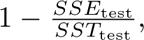 where *SSE*_test_ is the sum of squared prediction errors for the test set, and *SST*_test_ is the total sum of squares for the test set. We performed 10,000 row per column permutations for each split, ensuring that the same object permutations were maintained across the splits. We also estimated the noise ceiling, representing the maximum predictable variance, to determine the need for model or data improvements [13]. This approach provided a robust framework to validate the predictive performance of our computational model against behavioral data. As additional comparison models, we considered four NLP embeddings (Word2Vec [15], BERT[16] and CLAP[17] with no dimensionality reduction applied) to compare semantic learning in our audio-based semDNN with text-based learning. For Word2Vec, we computed a single semantic embedding for each sound by taking the average of the semantic embeddings for the name and verb sound descriptors. However, for BERT and CLAP, we directly obtained a single semantic embedding for each sound by estimating the semantic embedding for the name plus verb sentence. We also considered four pre-published categorical sound-to-event CNNs (Yamnet[3], VGGish[3], Kell[44], and CNN-14 from PANNs models[45]) along with variants of the semDNN network ( SemDNN_BERT_,SemDNN_CLAP_,SemDNN_Unbal_, SemDNN_NoTrain_).

## 3 Results

In this section, we present the results of our experiments evaluating the performance of models in predicting semantic relations between sounds and matching human behavior in an auditory cognitive task.

The semantic predictivity of the networks was evaluated considering both the internal SuperHardDrive Combo dataset (used for training the networks), and also considering the external datasets (FSD50k, US8k, ESC-50, and MSOS) which were not used for the training or potential subsequent fine-tuning. Figure 7 shows the pairwise comparisons of the averaged Ranking Score across all evaluation sounds from the internal and external datasets for the four tested models: CatDNN, SemDNN with Word2Vec, SemDNN with CLAP, and the model trained with BERT. Additionally, Figure 8 showcases the AMCSS comparison between the models. The graph depicts the average AMCSS or Ranking score on the two axes, with the intersection representing the corresponding metrics for each model. In the graph, points below the line of equality indicate that the model on the x-axis performs better on that dataset, and vice versa. SemDNN trained with Word2Vec emerges as having the best performance across all the comparisons, outperforming competing models in terms of both AMCSS and Ranking scores (see figure 9, for bar plots of average performance metrics across all datasets).

**Fig. 7.**
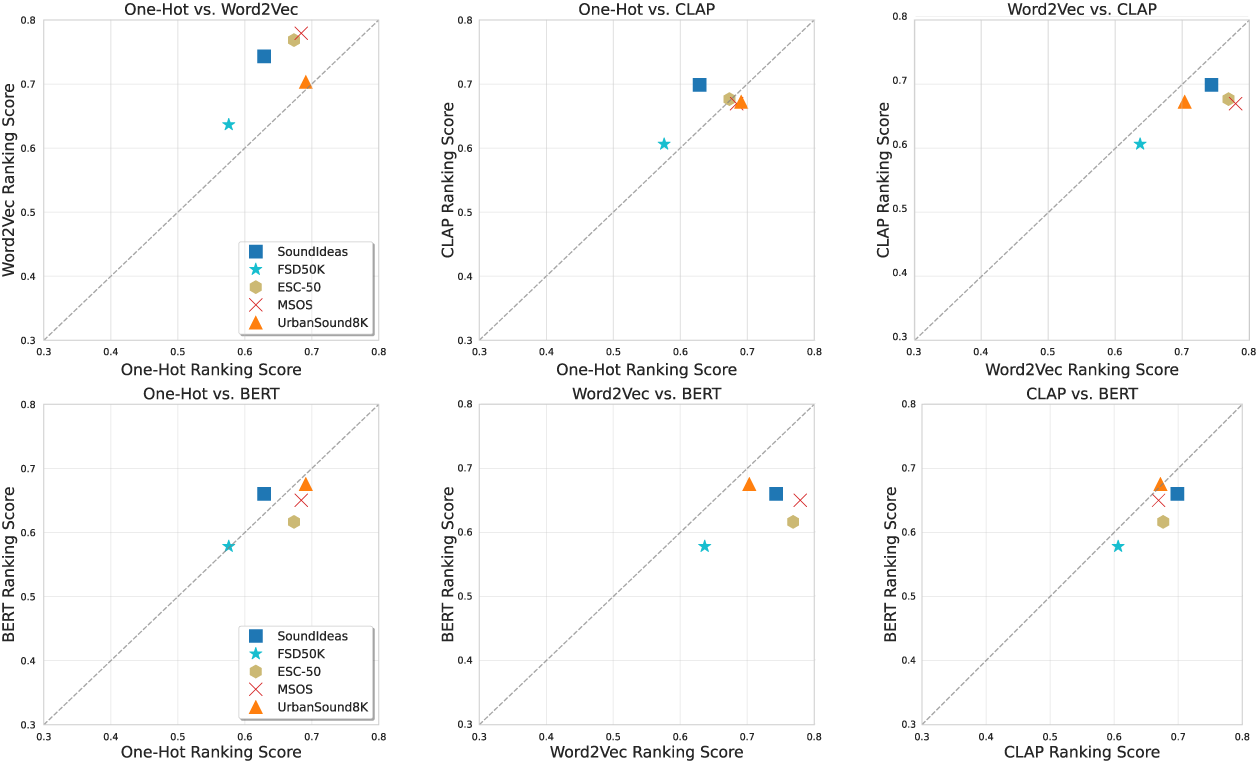
Pairwise Comparison of Ranking Scores Among Models. Averaged Ranking Scores of CatDNN, SemDNN with Word2Vec, SemDNN with CLAP, and SemDNN with BERT embeddings. Points below the equality line indicate better performance of the model on the x-axis for the corresponding dataset, and vice versa.

**Fig. 8.**
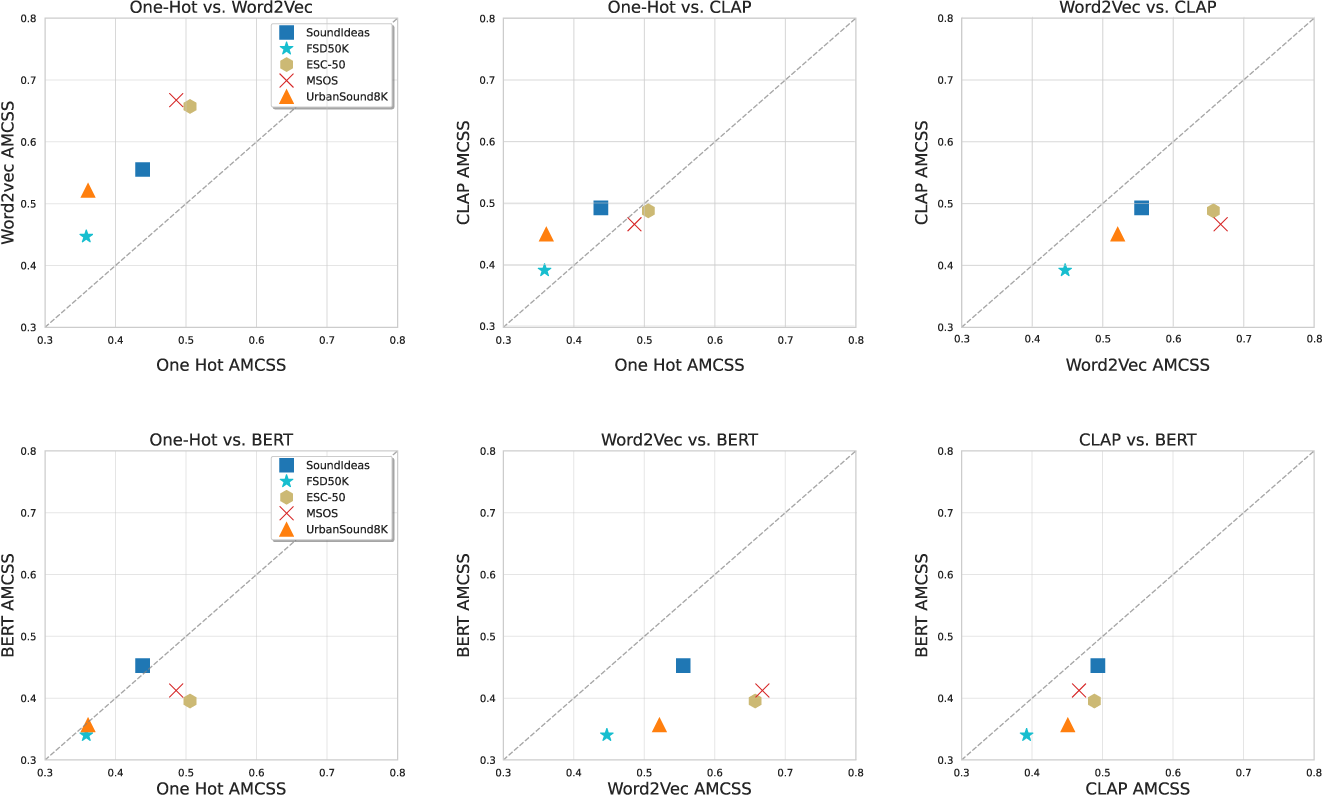
Pairwise Comparison of AMCSS Among Models. Average AMCSS (Average Maximum Cosine Similarity Score) for CatDNN, SemDNN with Word2Vec, SemDNN with CLAP, and SemDNN with BERT embeddings. Points below the equality line indicate superior performance of the model on the x-axis for the corresponding dataset, and vice versa.

**Fig. 9.**
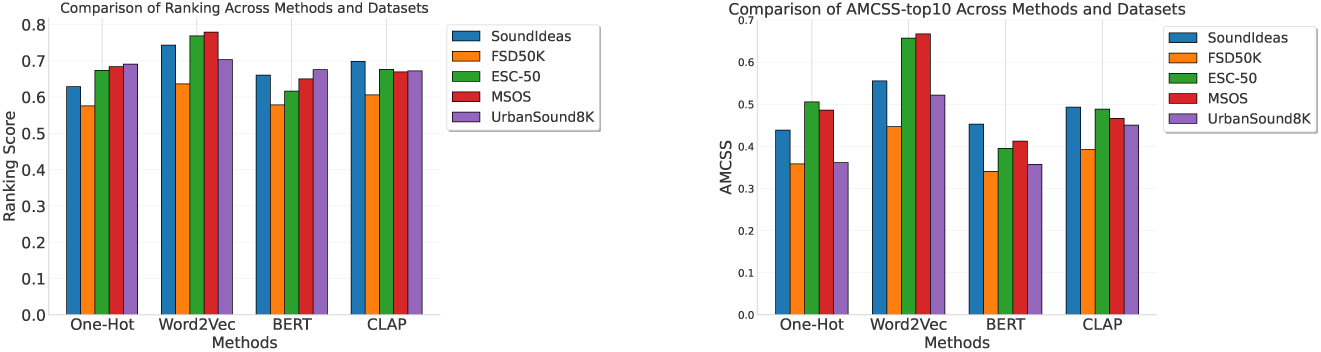
Bar plot summary comparison of Ranking Scores and AMCSS. The bar plot on the left represents the average Ranking Scores for CatDNN, SemDNN with Word2Vec, SemDNN with CLAP, and the BERT-trained model. The bar plot on the right represents the average AMCSS for the same models. SemDNN trained with Word2Vec consistently outperforms the other models.

Table 2 shows some examples of the top 5 predicted words retrieved from the NNLS, for SemDNN and its variants, and from the sigmoid activations, for catDNN. To show the results of our NLP pipeline (see section 2.3) in order to get labels from the SoundIdeas [43] dataset, we present in the first column the name of the filenames. Note in the last row how we moved from Harrier, which is a type of fighter jet, to its base-level category.

Our hypothesis was that SemDNN embeddings would outperform CatDNN in predicting higher-order semantic relations between sounds. To test this, we evaluated the MSOS dataset (see 2.4), where sounds are grouped into five macro-classes: sound effects, human, music, nature, and urban. For both SemDNN and CatDNN models, we computed pairwise normalized cosine distances between sound embeddings in the last intermediate layer (Fig. 2, arrow). In figure 10, the upper left panel illustrates an idealization, which synthetically reflects the original macro-class organization constructed in the left panel. In this synthetic construction, we assigned a within-category distance of 0 and a between-category distance of 1, allowing for a clear distinction between categories.

**Fig. 10.**
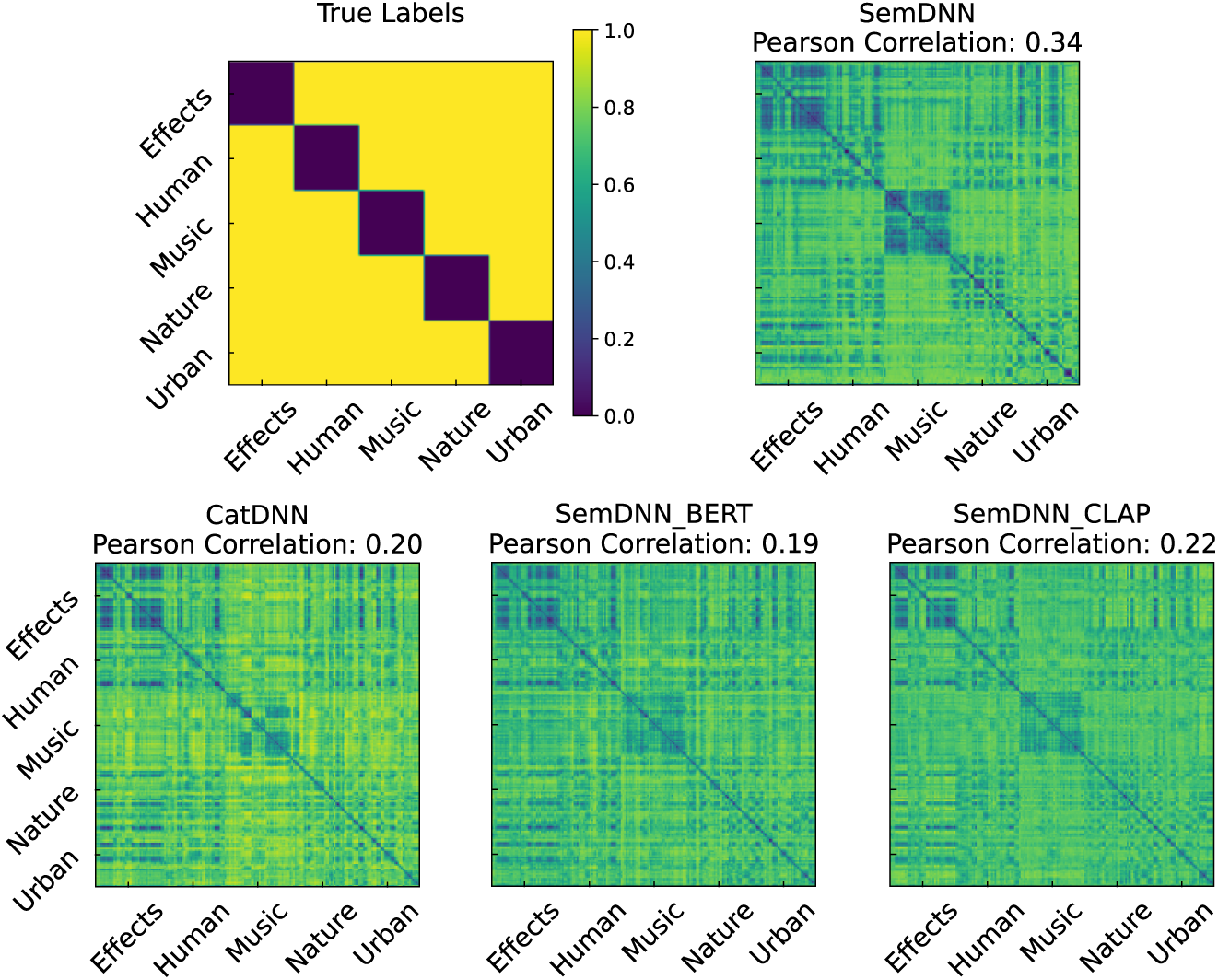
Comparison of Embedding Dissimilarity Matrices for SemDNNs and CatDNN. Normalized Cosine distances between MSOS sound embeddings of the interme-diate layer 512-D of CatDNN and SemDNN trained with Word2Vec, BERT, and CLAP representations (arrow in Figure 2). The matrix on the upper left reflects the true macro-classes, where the color scale represents the minimum (blue) and maximum (yellow) normalized distances, specifically within-category distance = 0; between-category distance = 1. The SemDNN trained with Word2Vec embedding matrix demonstrates a stronger reflec-tion of the macro-class organization compared to the CatDNN embedding matrix and the other two SemDNN variants, as indicated by higher Pearson correlation coefficients with the true categorical model: 0.340 for SemDNN with Word2Vec, 0.194 for SemDNN with BERT, 0.224 for SemDNN with CLAP and 0.193 for CatDNN.

To compare the performance of different SemDNN variants and CatDNN, we calculated normalized cosine distances and Pearson correlation coefficients between the synthetic matrix and the computed dissimilarities. The matrix in the upper right panel represents the SemDNN trained with Word2Vec embeddings dissimilarities, while the matrix in the lower left panel represents the CatDNN embeddings. The left two panels are the dissimilarities of SemDNN trained with BERT and CLAP, respectively. The color scale in all the matrices represents the normalized distances, ranging from minimum (blue) to maximum (yellow).

Among the SemDNN variants, the SemDNN model trained with Word2Vec embeddings demonstrated a stronger correlation with the true categorical model, with a Pearson correlation coefficient of 0.340. Comparatively, the SemDNN models trained with BERT and CLAP embeddings exhibited lower correlation coefficients of 0.194 and 0.224, respectively. The CatDNN embedding displayed the weakest correlation with the true categorical model, with a Pearson correlation coefficient of 0.193.

Figure 11 shows the ability of the various models to predict behavioral, sound, and word, dissimilarities using the cross-validated R-squared statistic. Models are grouped in four classes: semantics (blue), acoustics(light blue), CatDNNs(green), and SemDNNs (red). The noise ceiling represents the upper bound or best possible performance given data limitations or experimental constraints.

**Fig. 11.**
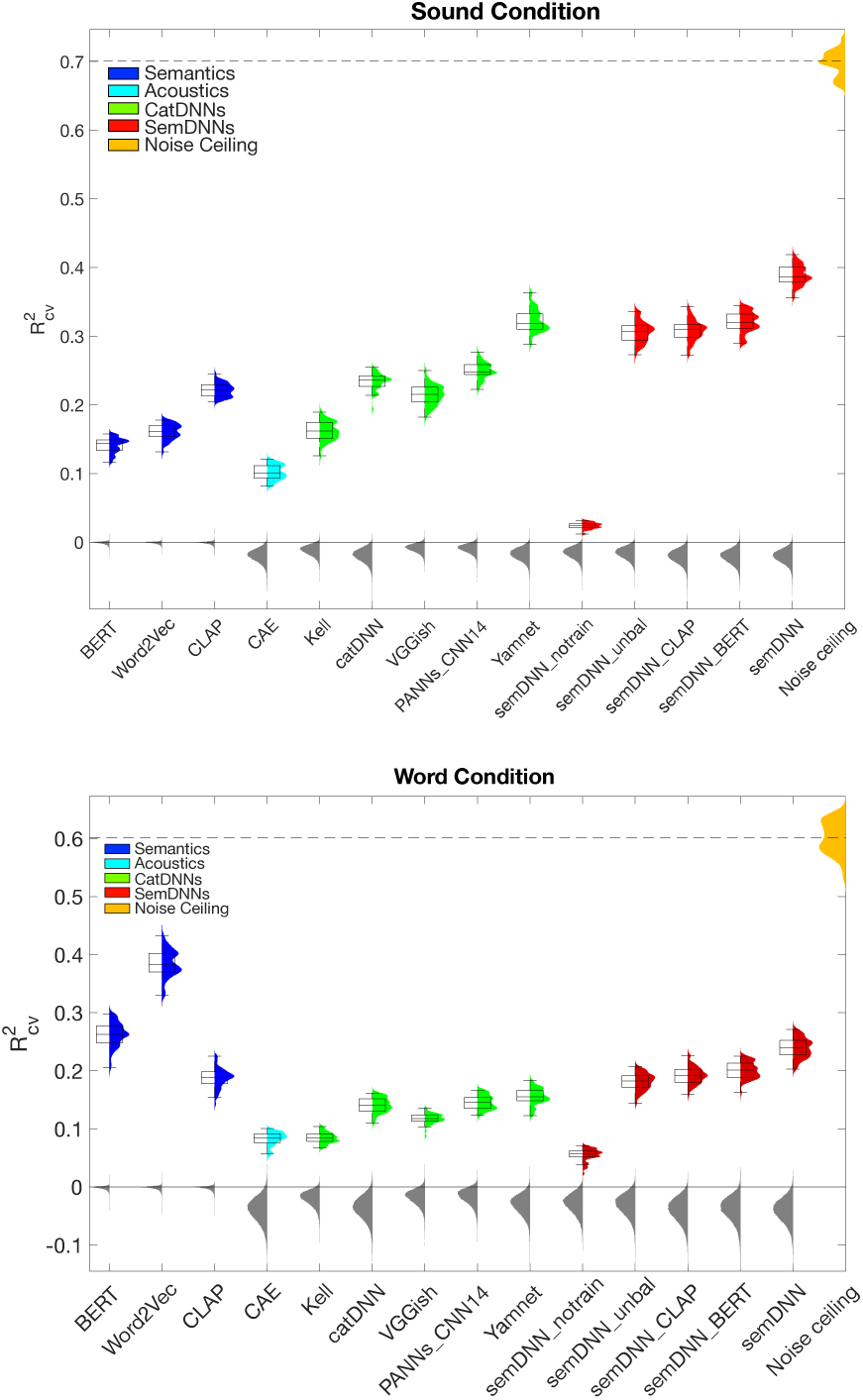
Cross-validated RSA results for sound condition and word condition. The color distributions correspond to the plug-in distribution of 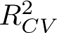 across CV folds, rep-resented by the box plots. The center of the box plot represents the median, while the lower and upper box limits indicate the 1st and 3rd quartiles, respectively. The bottom and top whiskers depict the data within 1.5 interquartile ranges from the 1st and 3rd quartiles, respectively. The dark gray color represents the cross-CV fold median of the permutation results. The orange color indicates the noise ceiling, with the dashed line representing the median noise ceiling across CV folds. The upper graph shows the performance of the evaluated models in predicting perceived sound dissimilarity (SemDNN outperforms all other models). The lower graph shows the performance of the evaluated models in perceived word dissimilarity, notably, Word2Vec outperforms all the other models.

Table 3 presents a summary of the two RSA results, highlighting, on the left column, SemDNN’s superior predictivity of perceived sound dissimilar-ity (highest 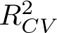 value) than CatDNN and other models (see also figure 11). Importantly, SemDNN outperformed all the competing networks trained with categorical labels (VGGish, PANNs CNN-14, Yamnet, and Kell). CLAP was instead the most predictive of the semantic models. These results confirm our hypothesis that a network that learns continuous semantic representations from acoustics better approximates human behavior compared to models con-sidering only acoustic information, relying solely on semantic information, or learning categorical semantic representations from acoustics. SemDNN trained with Word2Vec representations outperformed SemDNN trained with CLAP or BERT representations. Additionally, training SemDNN on a semantically bal-anced dataset yielded better results compared to training on a randomly chosen dataset (semDNN_unbal_), highlighting the importance of a balanced dataset [37]. We also evaluated the performance of an untrained network (semDNN_notrain_) initialized with random values, serving as a baseline. The right column of Table 3 focuses on the prediction of word perceived dissimilarity of the considered models measured by their respective 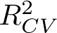 values. Notably, Word2Vec emerges as the most successful semantic model in this task. This outcome is in line with our expectations, considering the nature of our labels, which are keywords, and not actual sentences, representing the semantic content of the sounds. Among the DNN models, SemDNN stands out as the top performer in predicting word-perceived dissimilarity. This can be attributed to the fact that SemDNN is trained using Word2Vec representations, which aligns well with our keyword-based labels. The inherent strength of Word2Vec in capturing semantic relationships and similarities enables SemDNN to leverage this knowledge effectively, resulting in superior performance compared to other DNN models.

**Table 3.**
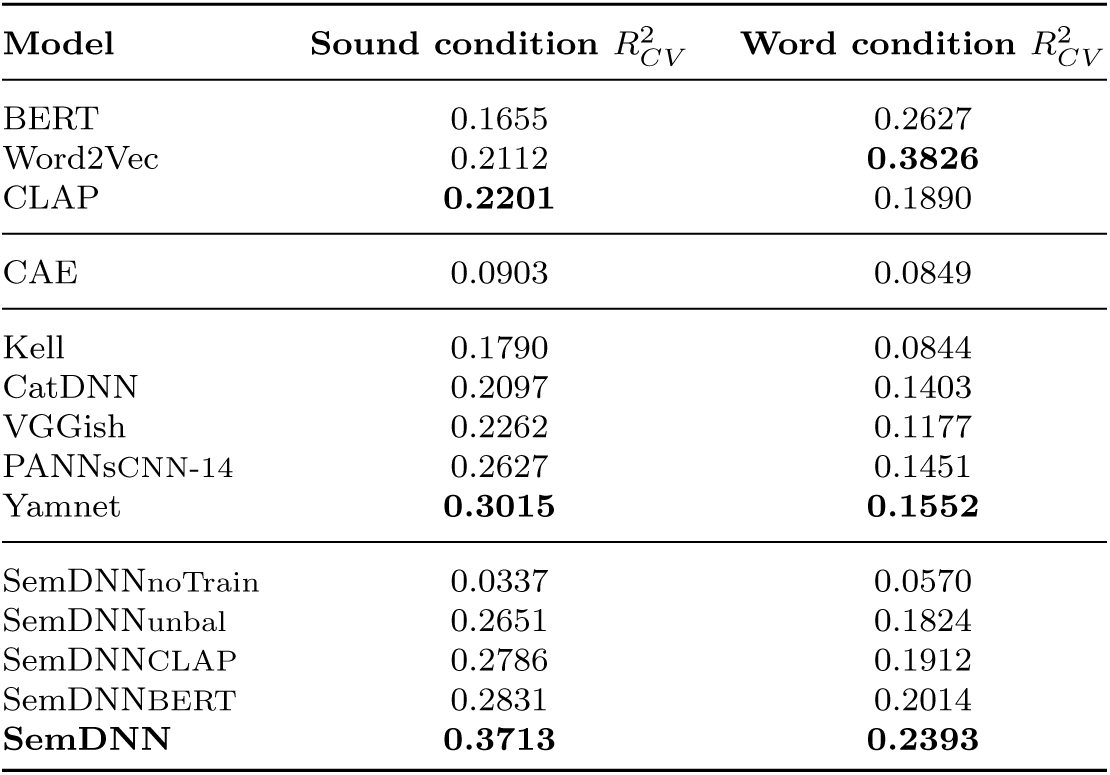
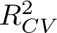obtained from the cross-validated RSA.

| Model | Sound condition $R_{CV}^2$ | Word condition $R_{CV}^2$ |
| --- | --- | --- |
| BERT | 0.1655 | 0.2627 |
| Word2Vec | 0.2112 | **0.3826** |
| CLAP | **0.2201** | 0.1890 |
| CAE | 0.0903 | 0.0849 |
| Kell | 0.1790 | 0.0844 |
| CatDNN | 0.2097 | 0.1403 |
| VGGish | 0.2262 | 0.1177 |
| PANNSCNN-14 | 0.2627 | 0.1451 |
| Yamnet | **0.3015** | **0.1552** |
| SemDNNnoTrain | 0.0337 | 0.0570 |
| SemDNNunbal | 0.2651 | 0.1824 |
| SemDNNCLAP | 0.2786 | 0.1912 |
| SemDNNBERT | 0.2831 | 0.2014 |
| **SemDNN** | **0.3713** | **0.2393** |

## 4 Discussion

We conducted a systematic exploration of the impact of employing continuous semantic embeddings (*Word2Vec, BERT, and CLAP*) in training DNNs for sound recognition, contrasting them with categorical labels (*one-hot encoding*).

Through our experiments and analyses, we gained significant perspectives into how the choice of semantic representations influences the performance of artificial hearing algorithms.

We compared the different models and the categorical model by using averaged Ranking Scores and AMCSSs (figures 7 and 8) on various datasets (FSD50k, US8k, ESC-50, MSOS, and the internal SuperHardDrive Combo dataset). The results consistently demonstrated that SemDNN trained with Word2Vec outperformed CatDNN and SemDNNs trained with CLAP and BERT. These findings imply that training DNNs to map sounds into a dense space preserving semantic relationships between sound sources enhances the network’s ability to recognize and comprehend *individual* sound events. The superiority of semDNN trained with Word2Vec over those using BERT and CLAP suggests that the complexity of the optimal semantic space lies between a categorical representation, lacking semantic relations, and a context-dependent natural language space, which may involve excessively fine-grained information. Our study employed keyword labels instead of full sentences, potentially limiting models’ contextual learning. While Word2Vec performs well with keywords, BERT and CLAP are both optimized for sentence-level context and might have faced limitations in this keyword-based setup. It may be interesting, in future work, to conduct similar analyses with sounds described with fully-formed sentences, such as those used in automated-captioning challenges [46]. Nonetheless, Word2Vec outperformed these models, suggesting that natural sound semantics may not require complex contextual information for comprehension. This finding challenges the traditional view of natural sound perception’s semantic complexity, often examined through the lens of language semantics [47]. It suggests that the inherent characteristics of natural sounds, well-captured by Word2Vec’s relatively simple semantic mapping, may not necessitate the contextual information demanded by language semantics.

Our hypothesis was that DNNs that are trained to recognize sounds and simultaneously learn the semantic relation between the sources would mimic human behavior better than other existing networks. We assessed this hypothesis in two steps: First, we examined the ability of the DNNs to form higher-order semantic classes; second, we assessed their ability to approximate human behavior in auditory cognitive tasks.

In the first step, we focused on the MSOS dataset, which organizes sounds into five macro-classes (effects, human, music, nature, and urban). We computed the pairwise cosine distances between sound embeddings in the last intermediate layer of SemDNN and CatDNN (figure 10). The results indicated that the SemDNN embedding better reflected the macro-class organization compared to the CatDNN embedding. The Pearson correlation coefficient with the true categorical model of the five macro-classes was higher for SemDNN (0.330) compared to CatDNN (0.193). Notably, semDNN (and the other networks) were not explicitly trained to group individual sounds into macro-classes. The superiority of semDNN over catDNN underscores that semDNNs leverage the semantic relation between sound sources in addition to the acoustic similarity of specific sounds.

To address the second question, we conducted a cross-validated RSA (Representational Similarity Analysis) and assessed the performance of different models in explaining perceived sound and word dissimilarity ratings (figure 11). The results, summarized in Table 3, demonstrated that for the sound condition, SemDNN achieved the highest 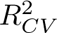 value among all the models, indicating its superior ability to emulate human behavioral data. It outperformed not only CatDNN and other DNNs trained with categorical labels but also purely semantic models such as Word2Vec, BERT, and CLAP. Interestingly, SemDNN trained with Word2Vec representations exhibited better performance than SemDNN trained with CLAP or BERT representations.

Moreover, we trained SemDNN with Word2Vec on a random choice dataset (SemDNN_unbal_) to compare the performance with a semantically balanced dataset. The results showed that SemDNN still outperformed SemDNN_unbal_, highlighting the importance of a semantically balanced dataset in training the model. Additionally, we included a baseline model (SemDNN_notrain_) that was untrained and solely initialized with random weights. For the word condition, the results are summarised in Table 3 and depicted in the lower graph of figure 11. Word2Vec outperforms other semantic models in predicting word dissimilarity, as expected. Since our labels are keywords rather than complete sentences, Word2Vec effectively captures the semantic content of the sounds. None of the CatDNNs stand out among the others showing the limitations of this network to perform a simple linguistic task. On the other hand, Word2Vec’s ability to capture semantic relationships and similarities enables SemDNN to leverage this knowledge effectively, leading to superior performance compared to other DNN models.

Overall, these results provide strong evidence for the effectiveness of SemDNN in capturing both acoustic and semantic information and approximate human behavior in auditory cognitive tasks. Integrating acoustic and semantic features proved more successful than considering acoustic information alone (CatDNN) or relying solely on pure semantic models. It’s worth noting that SemDNNs were trained with over 1 million examples, while 1 billion examples were considered to train VGGish and Yamnet [6], supporting the idea that - when the goal is to approxiamte human behaviour - ecological, balanced datasets may be more relevant than large amounts of unbalanced training data [37].

## 5 Conclusions

In this study, we investigated the performance of various models in performing sound recognition tasks and in their ability to approximate human behavior in auditory cognitive tasks (sound dissimilarity ratings). Our findings provide an important understanding of the role of semantic information in these two aspects. The key conclusions drawn from our analysis are as follows:

1. SemDNN, combining both acoustic and semantic information, consistently outperformed CatDNNs consistently outperformed CatDNNs in both sound recognition performance and approximating human behavioral data. This suggests that our approach of mapping sounds to a continuous space is a valid and advantageous alternative to the conventional method of training sound-to-event DNNs for discrete sound categories.
2. SemDNN models trained with Word2Vec representations exhibited superior performance compared to other semantic representations like BERT or CLAP. This underscores the effectiveness of Word2Vec embeddings in basic sound recognition tasks. Future work should explore the generalizability of these findings, especially when using datasets with complex linguistic descriptions of sounds.
3. Training SemDNN models on a semantically balanced dataset improved the prediction of human behavioral data compared to training on a randomly chosen dataset. It outperformed many other models trained on a larger number of sounds, emphasizing the importance of dataset curation and the use of ecologically valid datasets, particularly when aiming to approximate human behavior.

In summary, our study advances our understanding of the interplay between acoustics and semantics in both sound-to-event DNNs and human listeners. This paves the way for future research to optimize models and enhance their alignment with human perceptual judgments.

## Acknowledgments

Funding/Support: This work was supported by the Dutch Research Council (NWO 406.20.GO.030 to EF), the French National Research Agency (ANR-21-CE37-0027-01 to BLG; ANR-16-CONV-0002 – ILCB; ANR11-LABX-0036 – BLRI), Data Science Research Infrastructure (DSRI; Maastricht University) and the Dutch Province of Limburg.

## Data Availability

The data that support the findings of this study are available from SoundIdeas Inc. [43] but restrictions apply to the availability of these data, which were used under Royalty free license[48] for the current study, and so are not publicly available.

